# Tension promotes kinetochore-microtubule release in response to Aurora B activity

**DOI:** 10.1101/2020.06.01.127795

**Authors:** Geng-Yuan Chen, Fioranna Renda, Huaiying Zhang, Alper Gokden, Daniel Z. Wu, David M. Chenoweth, Alexey Khodjakov, Michael A. Lampson

## Abstract

Aurora B kinase regulates kinetochore-microtubule interactions to ensure accurate chromosome segregation in cell division. Tension provides a signal to discriminate attachment errors from bi-oriented kinetochores with sisters correctly attached to opposite spindle poles. Current models focus on tension as an input to locally regulate Aurora B activity. Here we show that the outcome of Aurora B activity depends on tension. Using an optogenetic approach to manipulate Aurora B at individual kinetochores, we find that kinase activity promotes microtubule release when tension is high. Conversely, when tension is low, Aurora B activity promotes depolymerization of kinetochore-microtubule bundles while maintaining attachment. We propose that tension is a signal inducing distinct error-correction mechanisms, with release or depolymerization advantageous for typical errors characterized by high or low tension, respectively.

## Main Text

To maintain genome integrity during cell division, kinetochores of sister chromatids attach to opposite spindle poles (bi-orientation) to ensure accurate segregation. This process relies on robust mechanisms to both identify errors by distinguishing correct from incorrect attachments and correct errors by changing the connections between kinetochores and microtubules. To identify errors, tension from opposite spindle poles is widely accepted as a signal indicating correctly bi-oriented sister kinetochores, whereas lack of tension signals an error (*1–4*). Correcting errors depends on destabilizing kinetochore-microtubule interactions so that new attachments can form. Aurora B kinase is a key regulator that destabilizes these interactions by phosphorylating kinetochore substrates that bind microtubules (*5–7*). To couple error identification and correction, current models propose that phosphorylation of Aurora B substrates depends on tension, through several possible mechanisms: separation of kinetochores from Aurora B at the inner centromere (*8–11*), regulation of Aurora B activity (*12*), or Aurora B localization to a kinetochore binding site (*13–15*).

An alternative model, which has not been tested, is that tension could regulate the downstream response to Aurora B substrate phosphorylation. Two mechanisms by which Aurora B destabilizes chromosome attachments have been proposed: release or depolymerization (*3*) (**Figure S1A**). The release model, in which kinetochores detach from microtubules, is supported by abundant biochemical evidence of Aurora B substrate phosphorylation lowering the kinetochore-microtubule affinity (*2, 6, 7*). Alternatively, cellular studies suggest that Aurora B induces depolymerization of kinetochore-attached microtubules, without full detachment. When sister kinetochores are attached to a single pole (syntelic attachment), depolymerization pulls both towards the pole (*16*), where they subsequently detach due to pole-localized activities (*17, 18*). To distinguish between the release and depolymerization models, we acutely recruited Aurora B to kinetochores using a photocaged small molecule that heterodimerizes HaloTag and *Escherichia coli* dihydrofolate reductase (eDHFR) fusion proteins (**Figure 1A, S1B**) (*19, 20*). We fused HaloTag to the kinetochore protein SPC25 and eDHFR to the INBox subdomain of INCENP, which recruits and activates Aurora B as part of the chromosome passenger complex (*6*), and thus can be used to control localized Aurora B activity (*11, 21*) (**Figure 1B**). The release and depolymerization models predict different outcomes of Aurora B activation. Microtubule release reduces or eliminates forces exerted by kinetochore fibers (K-fibers), whereas depolymerization increases force pulling the kinetochore towards the attached pole. Using kinetochore movement as a readout, we tested these predictions with both monopolar and bipolar spindles, representing low and high tension conditions, respectively.

**Figure. 1.**
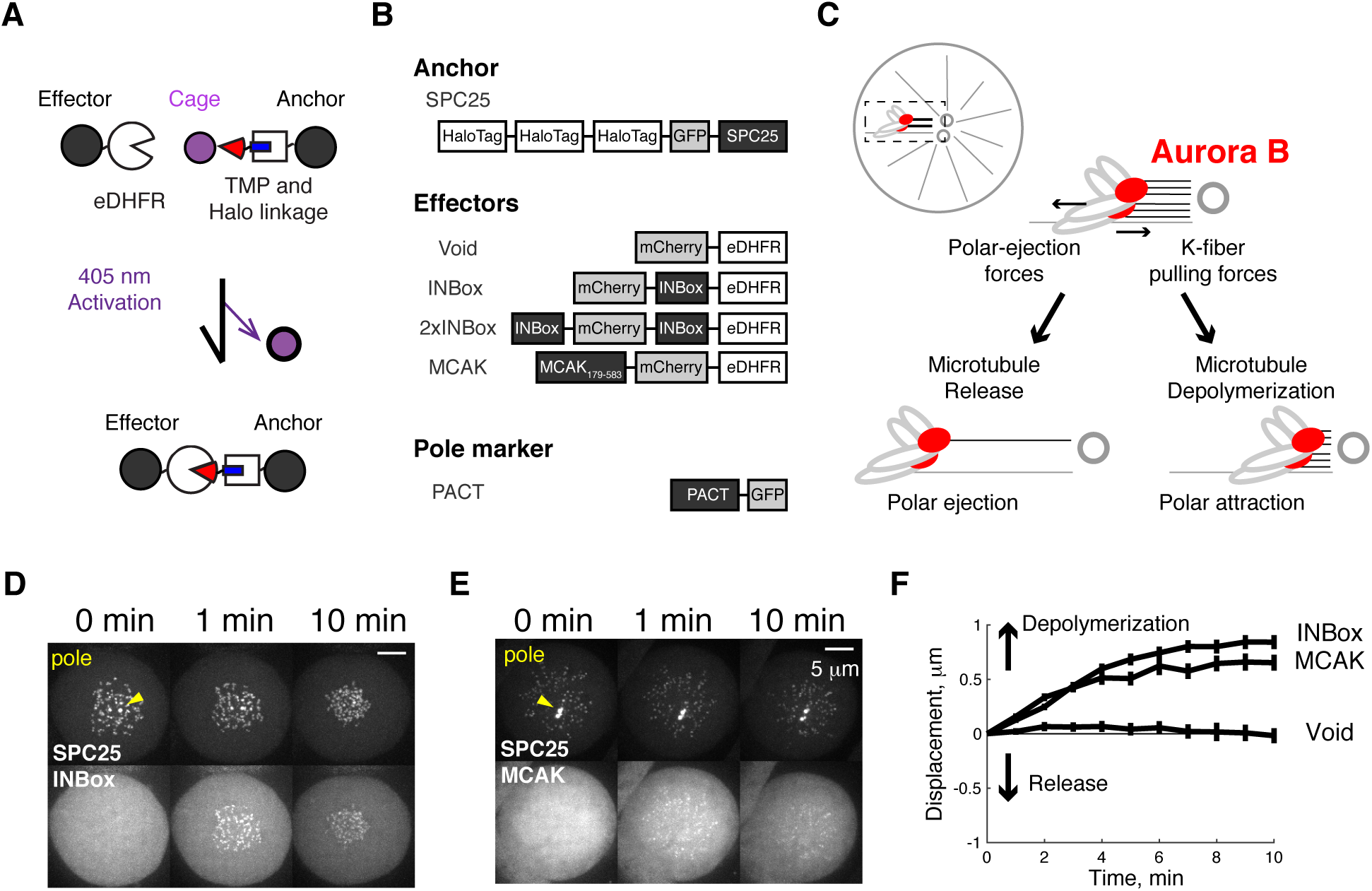
Recruitment of Aurora B to syntelic kinetochores triggers microtubule depolymerization. (**A**) Light-induced dimerization schematic. The dimerizer has a Halo ligand (blue) linked to the eDHFR ligand Trimethoprim (TMP, red), which is protected by a photo-activatable cage (purple). The kinetochore protein SPC25 anchors HaloTag at the outer kinetochore, and effectors are fused to eDHFR. Effectors are recruited to kinetochores by uncaging the HaloTag-bound dimerizer with light. (**B**) Constructs for this study. Dimerizer uncaging recruits eDHFR-tagged effector proteins (or Void as a negative control). The PACT domain targets to centrosomes to label spindle poles (yellow triangles) (*50*). (**C**) Schematic of monopolar spindle assay, with chromosomes under a tug-of-war between K-fiber depolymerization and polar-ejection forces. Red circles: Aurora B-activated kinetochores. After full or partial release of kinetochore microtubules, polar-ejection forces dominate and chromosomes move away from spindle poles. In contrast, depolymerization increases poleward forces. (**D-E**) Representative images before and after uncaging at t=0 (**Movie S1-2**). (**F**) Kinetochore displacement over time: poleward movement defined as positive direction, mean ± SEM (n ≥ 27 cells).

Inhibition of kinesin-5 (Eg5) with S-trityl-L-Cysteine (STLC) generates monopolar spindles, so that kinetochores are under low tension with frequent syntelic attachments (*22*). Normally, the combination of K-fiber dynamics and polar ejection forces dictates chromosome positions relative to the poles (**Figure 1C**). After Aurora B activation, the release model predicts chromosome movement away from the pole due to reduced forces from K-fibers, whereas the depolymerization model predicts movement towards the pole due to increased pulling forces from K-fibers. We found that INBox recruitment to kinetochores triggered poleward chromosome movement (**Figure 1D, F, Movie S1**), consistent with Aurora B inducing microtubule depolymerization as shown previously for syntelic attachments (*16*). As a positive control, we recruited the microtubule depolymerase MCAK (kinesin-13)(*23*) and observed similar poleward movement (**Figure 1E-F, Movie S2**).

Next, we examined the effects of Aurora B recruitment when sister kinetochores are bi-oriented on bipolar spindles and under high tension. We targeted individual kinetochores in this configuration because the release and depolymerization models make distinct predictions when a single kinetochore out of a pair is activated (**Figure 2A**). Microtubule release should lead to reduced pulling force from the activated kinetochore and movement away from the pole to which it was initially attached, whereas depolymerization has the opposite effect. We found that ∼50% of activated kinetochores moved beyond the bounds defined by the range of naturally-occurring chromosome oscillations (**Figure S2A-B, Movie S3**). We defined each of these events as depolymerization or release, denoted as positive or negative direction displacement, respectively. Upon INBox recruitment, release events were more frequent than depolymerization events (67% vs. 33%) (**Figure 2B, D, Movie S4**; **S2C-D, Movie S5**). Increasing Aurora B recruitment by using a construct with two copies of INBox (2xINBox, **Figure 1B**) further increased the proportion of release events to 87% (**Figure 2C, E, S2D, Movie S6**). After an initial lag phase, the released kinetochores moved with a steady-state velocity (∼1.5 μm/min) consistent with kinetochore ablation assays mimicking release (**Figure 2F, S2E**) (*24*). Furthermore, inter-kinetochore distances decreased after Aurora B recruitment (**Figure S2F**), consistent with relaxed tension predicted by microtubule release from one kinetochore. Together, these results indicate that bi-oriented kinetochores under tension primarily release their attached microtubules upon phosphorylation.

**Figure 2.**
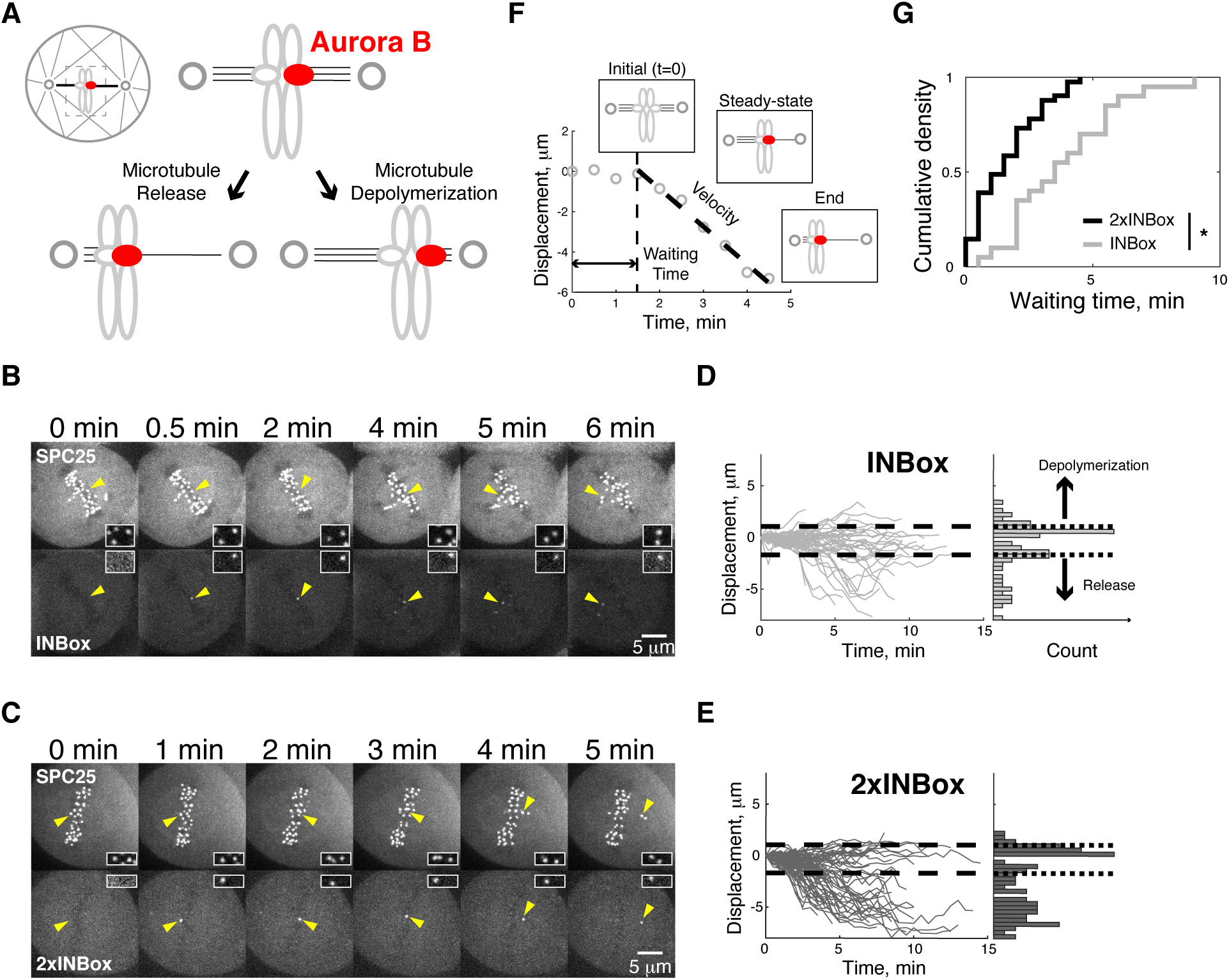
Recruitment of Aurora B to a single kinetochore of a bi-oriented pair triggers microtubule release. (**A**) Schematic of bipolar spindle assay. The bi-oriented sisters are under a tug-of-war between the two attached K-fibers. Microtubule release or depolymerization at the activated kinetochore (red circles) leads to movement in opposite directions. (**B-C**) Images from representative experiments showing INBox (B, **Movie S4**) or 2xINBox (C, **Movie S6**) recruitment after activation of a single kinetochore (yellow triangles) at t=0. Insets show the targeted kinetochore pair at higher magnification. (**D-E**) Displacement of the activated kinetochore from the metaphase plate over time: each trace represents a single kinetochore after INBox (D, n = 64) or 2xINBox (E, n = 79) recruitment, with the starting location defined as zero. Dashed lines show the range of chromosome dynamics covering 96% of control (Void-recruited) kinetochores (**Figure S2A-B, Movie S3**). Histograms show maximal displacement for each trace. (**F-G**) Analyses of released kinetochores after Inbox or 2xINBox recruitment. Example trace (F) shows waiting time after activation, followed by movement at steady-state velocity (**Figure S2E**). (G) Waiting time distribution (2xINBox: median = 1.5 min, n = 41; INBox: 3.5 min, n = 20). *P < 0.005.

To directly visualize microtubules after single kinetochore activation, we performed correlative serial-section electron microscopy (EM). Cells were fixed after the activated kinetochore moved out of the metaphase plate, indicating release (**Figure 3A**). We define the activated kinetochore as lagging and the sister kinetochore as leading because it moves towards its attached pole (**Figure 3B, B’**). We find microtubules bound to the leading kinetochore (Kb), but not to the activated lagging kinetochore (Ka) (**Figure 3B’, S3A’**). In contrast, kinetochore pairs that were not activated align properly on the metaphase plate, with K-fibers attached on both sides (**Figure S3A, A”**). Thus, the activated kinetochore releases its attached microtubules and thereby loses the tug-of-war to the sister. The coexistence of release and depolymerization events after Aurora B activation (**Figure 2B-E**) suggests an underlying kinetic race, with the release rate dominant on bipolar spindles. In addition, we find a shorter waiting time during the lag phase after recruitment of 2xINBox (1.5 min, median) vs. INBox (3.5 min), indicating that higher Aurora B activity further increases the microtubule release rate and the fraction of release events (**Figure 2G, S2D**). To explain the differences between monopolar and bipolar spindles, one possibility is that time in mitosis affects the outcome, as Eg5-inhibited cells are arrested in mitosis for up to 2-6 hours in our experiments. To test this possibility, we activated individual kinetochores in the presence of the APC/C inhibitor proTAME to delay mitotic exit in cells with bipolar spindles (**Figure S4A-B**). The fraction of release vs. depolymerization events and the release kinetics remain unchanged under these conditions (**Figure S4C-E**), indicating that time in mitosis is not a key variable.

**Figure 3.**
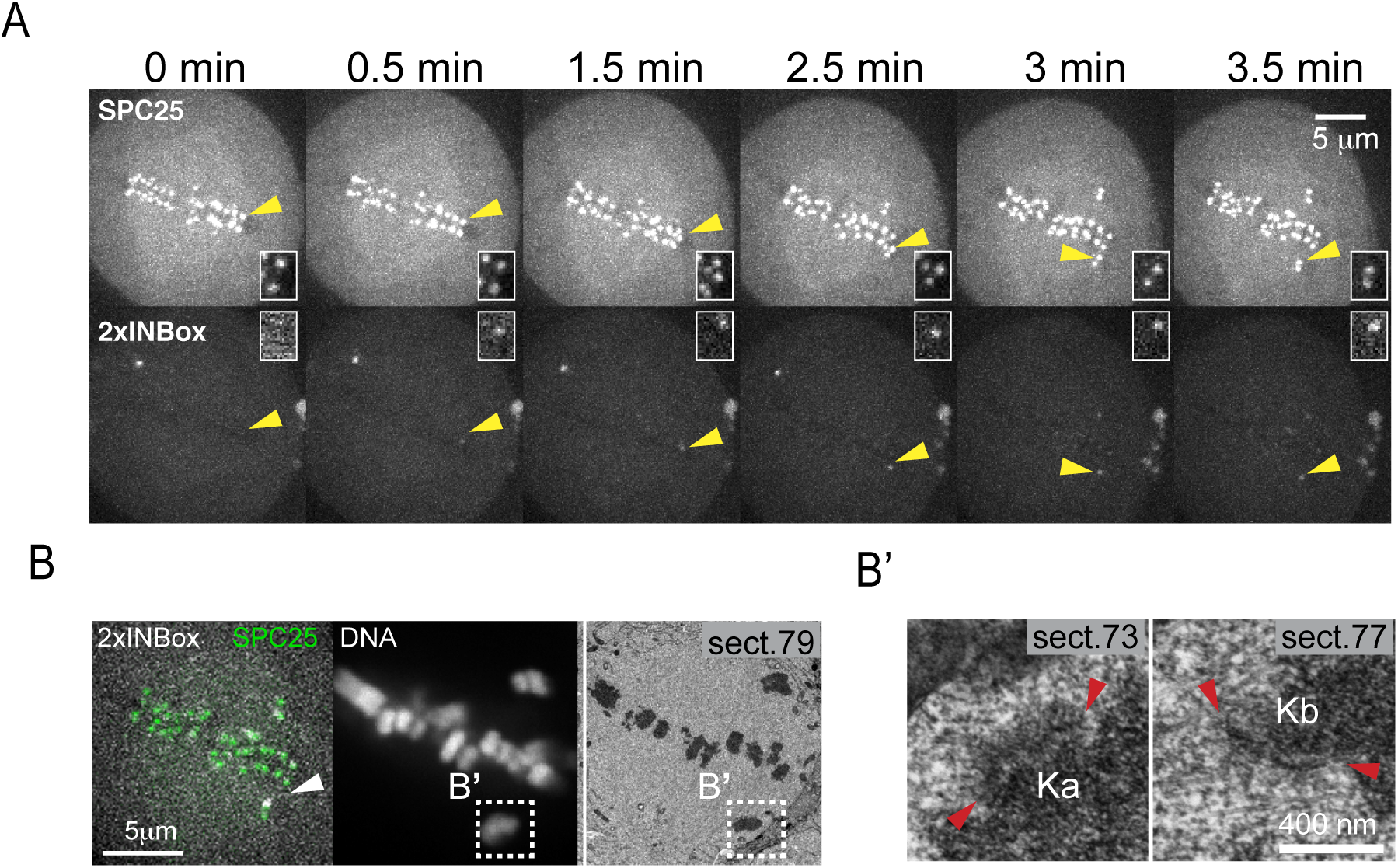
Activated kinetochores lack end-on microtubule attachments. (**A**) Live imaging showing 2xINBox recruitment and chromosome movement after activation of a single kinetochore (yellow triangles) at t=0, as in **Figure 2B**. (**B**) The same cell was fixed at 3.5 min and examined by fluorescence and serial-section EM. Left panels: a single focal plane shows 2xINBox overlaying SPC25 at the activated kinetochore (white arrow) and chromosomes by fluorescence, and the corresponding section from the 3D EM dataset. Kinetochores of the activated chromosome pair (boxes, **B’**) are shown at higher magnification in the right panels, with kinetochores bracketed by red triangles. Microtubules are bound to the leading kinetochore (Kb) but not the activated lagging kinetochore (Ka). Full EM series through these two kinetochores are shown in **Figure S3** together with serial sections from an aligned sister kinetochore pair that was not activated.

We next tested whether differences in tension, which is high on bipolar spindles but low on monopolar spindles, might explain the different outcomes in these two contexts. If tension promotes microtubule release upon Aurora B activation, then we predict that experimentally reducing tension on bipolar spindles would inhibit microtubule release. Reducing microtubule crosslinkers leads to ∼10% decrease in inter-kinetochore distance (*25, 26*), suggesting that interpolar microtubule arrays are mechanically coupled to K-fibers. We therefore tested whether the microtubule-crosslinking motors Eg5 or KIF15 (kinesin-12) mediate inter-kinetochore tension. Eg5 motors cross-link and slide microtubules apart to maintain proper pole-to-pole distance (*27–29*), while KIF15 motors additionally cross-link K-fibers to control chromosome movement (*30, 31*). Blocking nucleotide-binding of these motors entraps them in a rigor state, which maintains cross-linking activity but diminishes the powerstroke (*25, 32, 33*). We found that both the Eg5 rigor inhibitor BRD9876 and the KIF15 rigor inhibitor KIF15-IN-1 reduce inter-kinetochore distances by ∼10%, indicating reduced tension (**Figure 4A, S5A-B**). These inhibitors stabilize microtubules against depolymerization (*34, 35*) and therefore did not increase the fraction of depolymerization events after single kinetochore activation (**Figure S5C**). Addition of either inhibitor did, however, increase the median waiting time from 1.5 to 2.5 min for kinetochore release (**Figure 4B-D**), indicating that reduced tension slows the Aurora B-induced microtubule release rate.

**Figure. 4.**
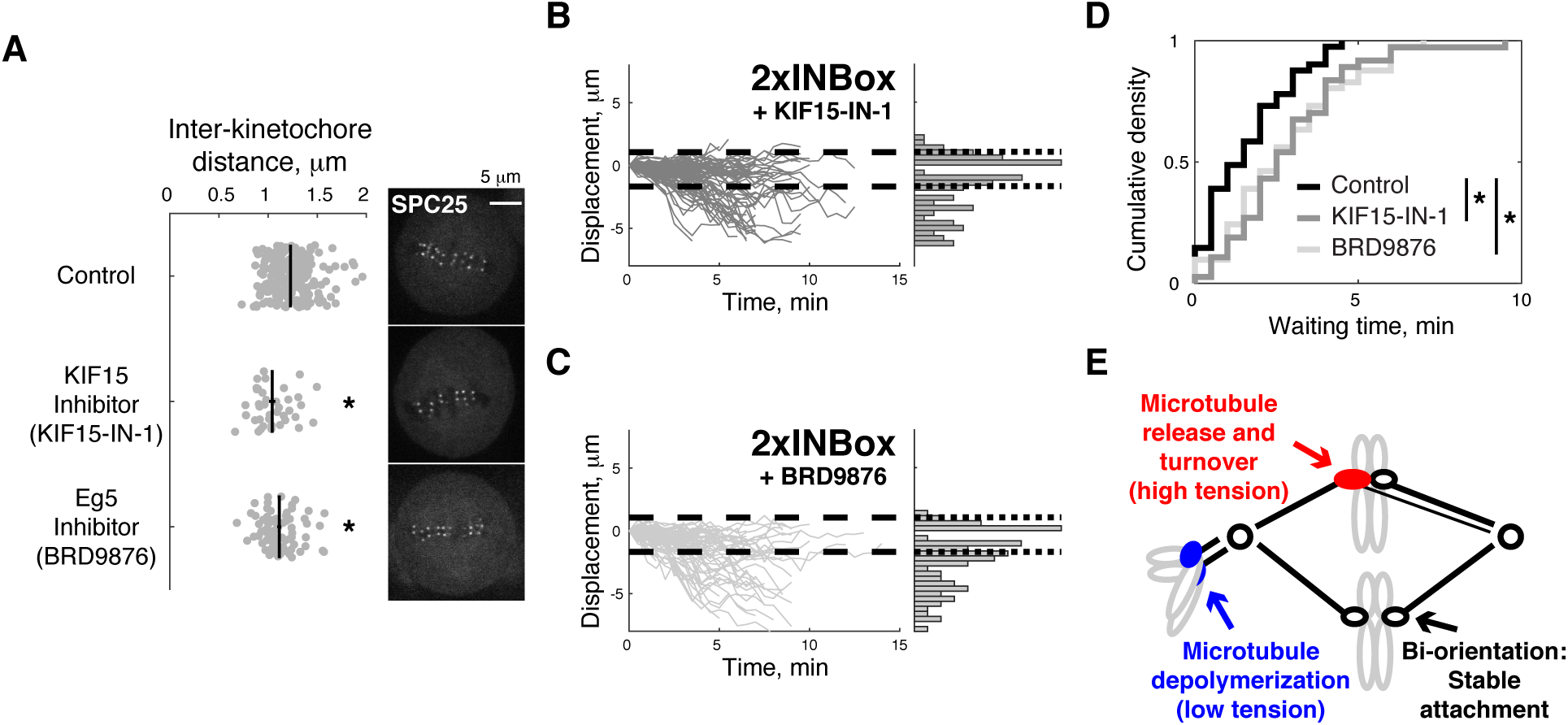
Reducing inter-kinetochore tension slows release of kinetochore microtubules. (**A**) Representative images of cells treated with a KIF15 inhibitor (40 μM KIF15-IN-1), Eg5 inhibitor (20 μM BRD9876), or control (no inhibitor). In the plot, each data point represents multiple kinetochores from a single cell. Black lines: mean ± SEM (n ≥ 40 kinetochores for each condition). (**B, C**) Displacement of the activated kinetochore after 2xINBox recruitment with KIF15 (B, n = 93) or Eg5 (C, n = 74) inhibition, plotted as in **Figure 2D**. (**D**) Waiting time distribution of the released kinetochores (control: median = 1.5 min, n = 41; KIF15-IN-1: 2.5 min, n = 37; BRD9876: 2.5 min, n = 41). *P < 0.05. (**E**) Model for distinct error-correction mechanisms at merotelic (red) and syntelic (blue) attachments in response to Aurora B activation.

Overall, our results demonstrate contrasting responses to Aurora B activity: depolymerization of kinetochore microtubules under low tension and release under high tension. This increase of the release rate with tension is opposite to the catch-bond behavior of unphosphorylated kinetochores, in which force stabilizes attachments (*36*). These seemingly contradictory effects of tension can be explained by a two-state model, in which the detachment rate is lower for polymerizing compared to depolymerizing microtubule plus-ends as shown *in vitro* (*2, 36*). At unphosphorylated kinetochores, the catch bond is due to tension promoting the polymerizing state, which is more strongly attached (*36*). Kinetochore substrate phosphorylation promotes microtubule depolymerization (*37*), preventing the transition to the more strongly attached state, which converts the catch bond into a more conventional slip bond that breaks under increasing force. Thus, phosphorylated kinetochores maintain attachment while depolymerizing under low tension, or release under higher tension. Phosphorylation also increases the detachment rate for polymerizing microtubules (*38*), and there may be additional complexity in the cell if phosphorylation of distinct Aurora B substrates (e.g., the Ndc80 or Ska complexes or MCAK) depends on tension, promoting either microtubule depolymerization or release (*6, 7*). Furthermore, centromeric microtubule depolymerases might confer distinct functions under the control of Aurora B (*39, 40*): drive poleward chromosome movement or generate tension at syntelic or bi-oriented attachments, respectively.

By integrating spindle mechanics and kinetochore biochemistry, we propose that the tension-dependent response facilitates correction of distinct attachment errors (**Figure 4E**). Lower tension is associated with increased Aurora B activity at syntelic attachments (*8, 41*), which typically scatter around spindle poles. Microtubule release would leave chromosomes to be pushed away from the spindle by polar ejection forces, whereas microtubule depolymerization pulls chromosomes towards the spindle, where they subsequently detach near the pole and congress by gliding along kinetochore fibers of other chromosomes (*17, 18, 42*). Aurora B is also activated at merotelic attachments, in which one kinetochore attaches to both spindle poles, either by recruitment or by interactions with microtubules (*43–45*). Because tension is higher with microtubules pulling in opposite directions, phosphorylation would promote microtubule release and turnover to achieve bi-orientation (*39, 40, 46*). This mechanism can explain the failure to correct merotelic errors associated with defective spindle mechanics, such as disruptions of poleward microtubule flux, tissue architecture, or tubulin homeostasis (*47–49*).

## Acknowledgments

We thank Dr. Ekaterina Grishchuk for commenting on the manuscript, other Philly Chromo Club members (Dr. Ben Black, Dr. Roger Greenberg, and Dr. Matthew Good) for constructive criticism, and the Physical Sciences Oncology Center and Penn Center for Genome Integrity for discussions.

## Author contributions

G.Y.C. designed and conducted cell biology experiments and wrote the manuscript. H.Z. and A.G. tested the feasibility of this project in the initial stage. F.R. and A.K. acquired and analyzed EM images. D.W. and D.M.C. synthesized and characterized the CTH dimerizer. M.A.L. designed experiments and edited the manuscript.

## Data and materials availability

All research resources, including protocols, constructs, cell lines, chemical probes, and MATLAB codes, are available upon request. The corresponding authors adhere to the NIH Grants Policy and Sharing of Unique Research Resources.

## Supplementary Materials for

### Materials and Methods

#### Photo-caged dimerizer

CTH synthesis and characterization followed published protocols (*20, 51*). Stocks were dissolved in DMSO at 20 mM and distributed in amber-colored 1.5 mL microcentrifuge tubes at −80 °C for long-term storage. A stock aliquot was diluted in growth medium to 10 μM working concentration with 1 mL volume and then kept at −80 °C until use.

#### Plasmids

All constructs were integrated into the pEM705 backbone containing a CAG promoter for constitutive expression (*20, 52*). The kinetochore protein SPC25 was chosen as an anchor because of its outer kinetochore localization and slow cytosolic exchange. The N-terminus of SPC25 was fused to 3 tandem copies of HaloTag and to GFP to make 3xHalo-GFP-SPC25. To recruit and activate Aurora B kinase, the INBox domain (human INCENP_819-918_) was fused to mCherry at its N-terminus and to *E. coli* DHFR (eDHFR) at its C-terminus to make mCherry-INBox-eDHFR. To maximize Aurora B activity, a second INBox domain was fused to the N-terminus of mCherry-INBox-eDHFR to make INBox-mCherry-INBox-eDHFR (2xINBox). As a negative control, mCherry was fused to the N-terminus of eDHFR to make mCherry-eDHFR (Void). As a positive control, human MCAK_179-583_ (*53, 54*) was fused to the N-terminus of mCherry-eDHFR to make MCAK-mCherry-eDHFR.

#### Cell cultures and transfection

All assays were carried out with HeLa RMCE acceptor cells stably expressing 3xHalo-GFP-SPC25, as previously described (*19, 20, 52*). Cells were cultured at 37 °C in growth medium containing Dulbecco’s Modified Eagle’s Medium plus 10% FBS and 1% penicillin-streptomycin, with 5% CO_2_ and a humidified atmosphere. For bipolar spindle assays, cells were grown on 22×22mm coverslips (Fisher Scientific) coated by poly-D-lysine (Sigma-Aldrich) for at least 16 hours, and then transfected with Lipofectamine 2000 (Invitrogen) plus 1-2 μg eDHFR-tagged constructs (Void, INBox, or 2xINBox) for another 24 hours. For monopolar spindle assays, 250 ng of PACT-GFP plasmid was co-transfected with the eDHFR-tagged constructs (Void, INBox, 2xINBox, or MCAK) for 24-36 hours.

#### Small-molecule inhibitors

Stocks were prepared in DMSO and stored at −20 °C. Stock concentrations were 10 mM S-Trityl-L-Cysteine (STLC, Eg5 inhibitor, Sigma Aldrich), 50 mM BRD9876 (Eg5 rigor inhibitor, Tocris), 20 mM proTAME (APC/C inhibitor, Boston Biochem), 20 mM KIF15-IN-1 (KIF15 rigor inhibitor, APExBIO). Fresh aliquots were used for all experiments.

#### Photo-activation and dimerization

Cells were incubated with CTH for 1h, followed by a 30-minute washout with growth medium to remove unbound molecules, as previously described (*20*). For imaging, coverslips were mounted in a magnetic chamber (Chamlide CM-S22-1, LCI) with L-15 medium without phenol red (Invitrogen) containing 10% FBS and 1% penicillin/streptomycin, then placed on a heated stage in a 37 °C environmental chamber (Incubator BL; PeCon GmbH). Targeted uncaging was performed using a 405 nm laser (CrystaLaser LC, model # DL405-050-O, 27 mW output after fiber coupling) under the control of an iLas2 software module (Roper Scientific) within MetaMorph (Molecular Device). To uncage CTH entirely, 8% laser power and 50 repetitions were used for monopolar spindle assays. For single kinetochore targeting, 7% laser power with 50 repetitions was used in a 590 nm diameter region. The region size was chosen to minimize off-target activation, based on the diffraction limit, the motion of kinetochores, and the time lag between imaging and manual activation.

For monopolar spindle assays or metaphase-arrested bipolar spindle assays, 20 μM STLC (*55*) or 20 μM proTAME (*56*) were added during CTH incubation and imaging. Kinesin rigor inhibitors were added by media exchange directly on the stage, followed by a 30-minute incubation to equilibrate the system. To avoid unwanted CTH uncaging during preparation stage, care was taken to minimize light and heat exposure, using low-luminescence or red light in the room and a long-pass filter to find cells under differential interference contrast microscopy.

#### Image acquisition

Live imaging was carried out with a confocal microscope (DM4000; Leica), equipped with a 100× 1.4 NA objective (Leica), an XY Piezo-Z stage (Applied Scientific Instrumentation), a spinning disk (Yokogawa), an electron multiplier charge-coupled device camera (ImageEM; Hamamatsu Photonics), and a laser merge module (LMM5; Spectral Applied Research) equipped with 488- and 593-nm lasers, as previously described (*20*). To minimize photo-bleaching in monopolar spindle assays, images were acquired with a 1 min time interval, with 1 μm spacing for GFP and mCherry z-stacks covering 15 μm total. To improve precision for the bipolar spindle assays, the time interval was 30 s with 0.5 μm z-spacing covering 3 μm total, so not all kinetochore pairs are visualized.

#### Image Processing and Data analyses

Images are shown as maximum-intensity z-projections. The Fiji plug-in, TrackMate, was used to define kinetochore coordinates globally. For monopolar spindle assays, the identified objects in the targeted cell were separated as kinetochores and poles based on their intensity and quality. Kinetochore and pole coordinates were imported into MATLAB (MathWorks). The distance from each kinetochore to the center of the monopolar spindle, defined by averaging the pole coordinates, was calculate to determine the average distance at each time point. Poleward movement was assigned as positive direction.

For bipolar spindle assays, the position of the metaphase plate was determined by fitting the interior kinetochore ensemble using linear regression, minimizing the mean-squares of x- and y-deviation, and kinetochores of unaligned chromosomes were omitted. The Fiji plug-in, MtrackJ, was used to manually track the activated single kinetochores and identify its sister kinetochore. The relative distances between the activated kinetochore and the fitted metaphase plate were calculated in MATLAB. Movement towards the pole attached to the sister kinetochore was defined as the negative direction. Hence, the depolymerization or release models predicts positive or negative directional movements, respectively, for both monopolar and bipolar spindle assays. The steady-state period of a released kinetochore was manually defined by the best linear fit of each trajectory moving beyond the negative threshold (an example in **Figure 2F**). Experiments were repeated at least three times on different days.

#### Correlative Electron Microscopy

Cells were fixed for 30 minutes in PBS containing 2.5% glutaraldehyde (Sigma) immediately after the last frame of live imaging. Complete Z-series at 0.2 µm steps were then recorded to map positions of chromosomes in Differential Interference Contrast (DIC) and Hoechst 33342 fluorescence (0.1 μg/ml, Life Technologies). The images were acquired on a Nikon eclipse Ti2E microscope with a Plan Apochromat 100×1.45 NA objective lens, Photometrics 95B Prime camera at 43-nm XY pixels. EM embedding, re-location of cells, and serial sectioning were done as previously described (*57*). 80-nm sections were imaged on a JEM 1400 microscope (JEOL) operated at 80 kV using a side-mounted 4.0 Megapixel XR401 sCMOS AMT camera (AMT). Complete images series recorded at 10K magnification were used to reconstruct partial volumes containing adjacent activated chromosomes. These volumes were aligned with the light microscopy images by matching positions of prominent landmarks such as chromosome arms. Serial higher-magnification images (40K) were then collected to detail distribution of microtubule in the vicinity of activated kinetochores. In two cells we successfully activated a single kinetochore, identified the activated kinetochore based on 2xINbox fluorescence signal in fixed cells, and analyzed it by correlative EM. In both cases, the activated kinetochore lacked end-on microtubule attachments as shown in the representative sample in **Figure 3**.

#### Statistical tests

A non-parametric Kolmogorov-Smirnov test was used to compare the cumulative waiting time distributions, and a two-sample Chi-square test was used to compare the fractions of release vs. depolymerization events. Titration curves were fit by a rectangular hyperbola to measure the IC_50_ of inter-kinetochore distance for KIF15-IN-1 and BRD9876. The two-sample velocity or inter-kinetochore distance comparisons were carried out using the Student’s t-test.

**Figure S1.**
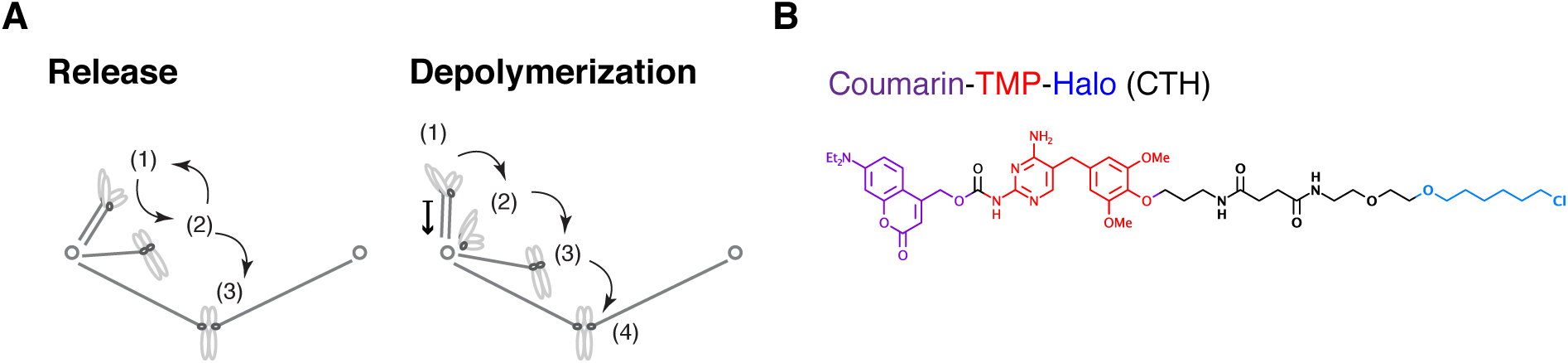
**Related to Figure 1**. (**A**) Two proposed models of error correction mediated by Aurora B. State 1 is the initial incorrect state. In the release model, microtubules detach from kinetochores (state 2) to allow new attachments to form. The process is iterative until bi-oriented attachments (state 3) are stabilized. In the depolymerization model, microtubules depolymerize while maintaining kinetochore attachment, generating poleward chromosome movement. Microtubules detach near the pole (state 2), followed by translocation along another K-fiber to the spindle equator (state 3), and binding of a microtubule from the opposite pole to achieve bi-orientation (state 4). (**B**) The photo-caged chemical dimerizer used in this study. CTH consists of the following components: a coumarin photocage; trimethoprim (TMP), which binds eDHFR; and a Halo ligand, which binds covalently to the HaloTag protein.

**Figure S2.**
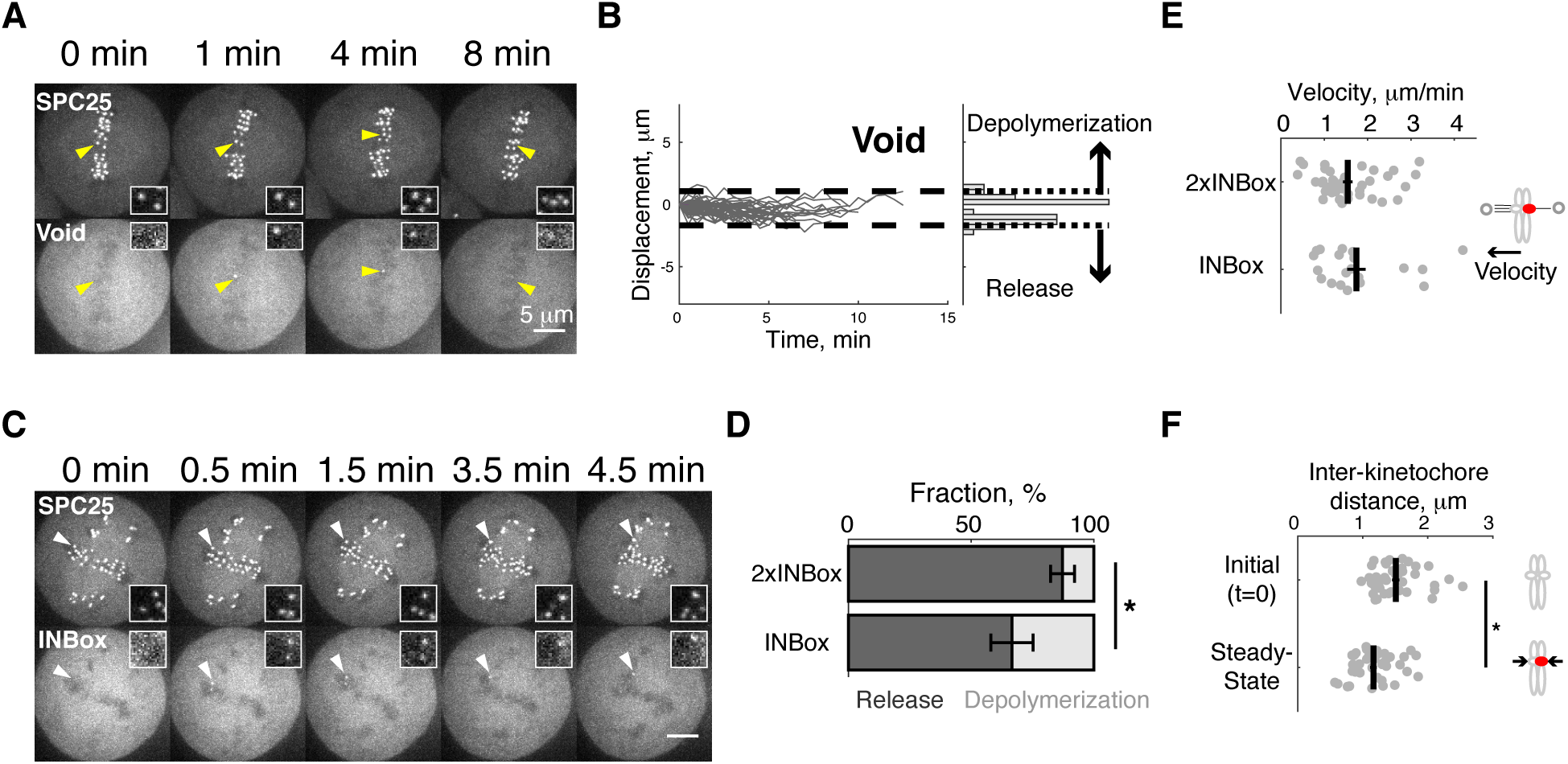
**Related to Figure 2**. (**A-B**) Representative images (A) and displacements (B) for Void recruitment to a single kinetochore of a bi-oriented pair, as a negative control. Displacements, plotted as in **Figure 2D-E**, were used to define the range of naturally-occurring chromosome oscillations (dashed lines cover 45 out of 47 traces, 96%). (**C**) Representative images showing kinetochore movement indicating microtubule depolymerization after recruiting INBox (**Movie S5**). Insets (A and C) show the targeted kinetochore pair at higher magnification. (**D**) The fraction of release vs. depolymerization events from **Figure 2D**-**E** (2xINBox: 87 ± 5% mean ± SEM, n = 47; INBox: 67 ± 9%, n = 30; *P < 0.05). (**E-F**) Steady-state velocities (E) and inter-kinetochore distances (F) for released kinetochores. Distances are plotted before activation and at steady-state after 2xINBox recruitment. Each data point represents a single kinetochore or pair of sisters, black lines: mean ± SEM (n = 41 for 2xINBox, n = 20 for INBox).

**Figure S3.**
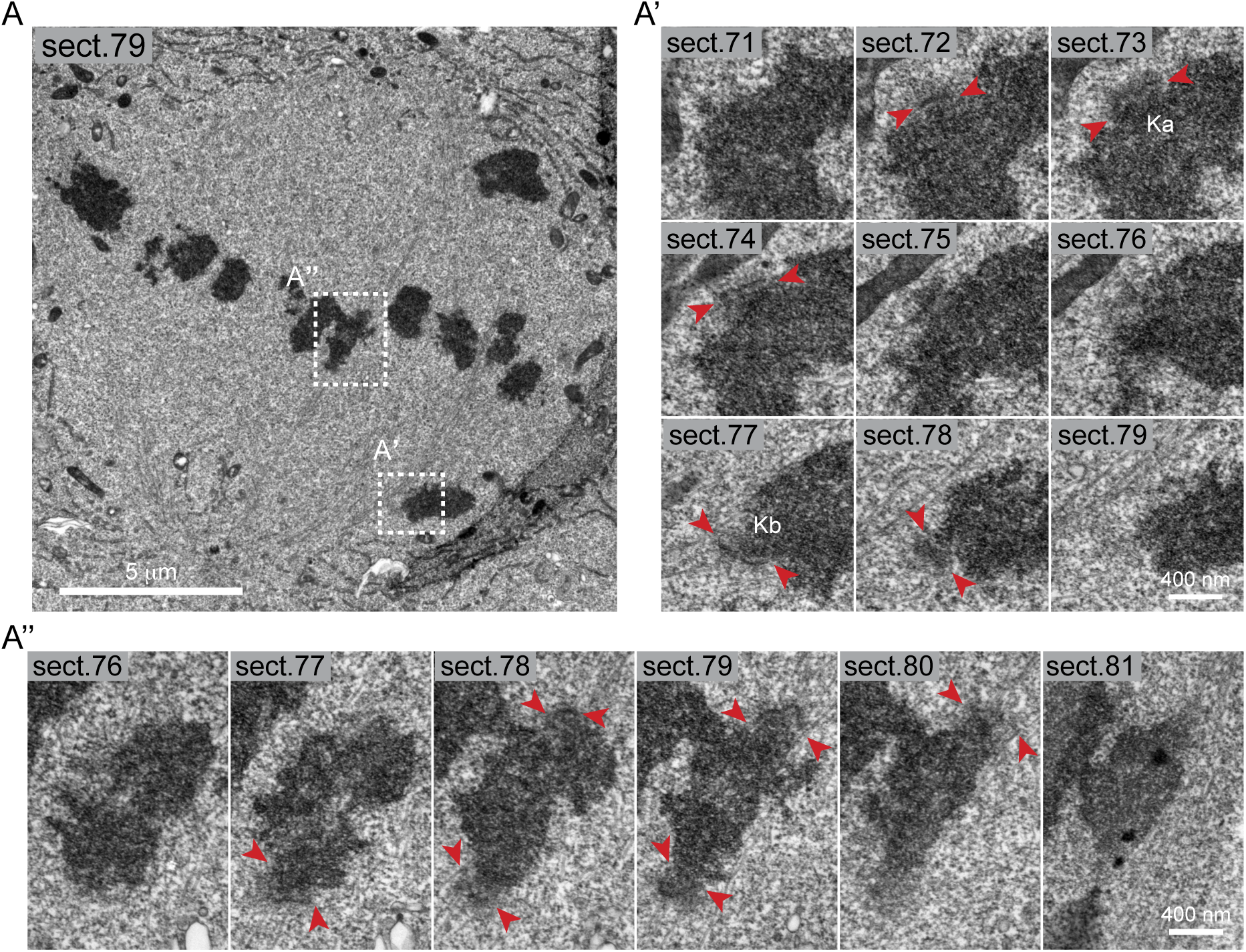
**Related to Figure 3.** (**A**) The magnified image from **Figure 3B**. A chromosome containing the activated kinetochore (**A’**) and a metaphase-aligned chromosome (**A”**) are boxed for a detailed 3D analysis. In contrast to the activated chromosome (**A’**), the aligned chromosome (**A”**) has roughly the same microtubule numbers on both kinetochores.

**Figure S4.**
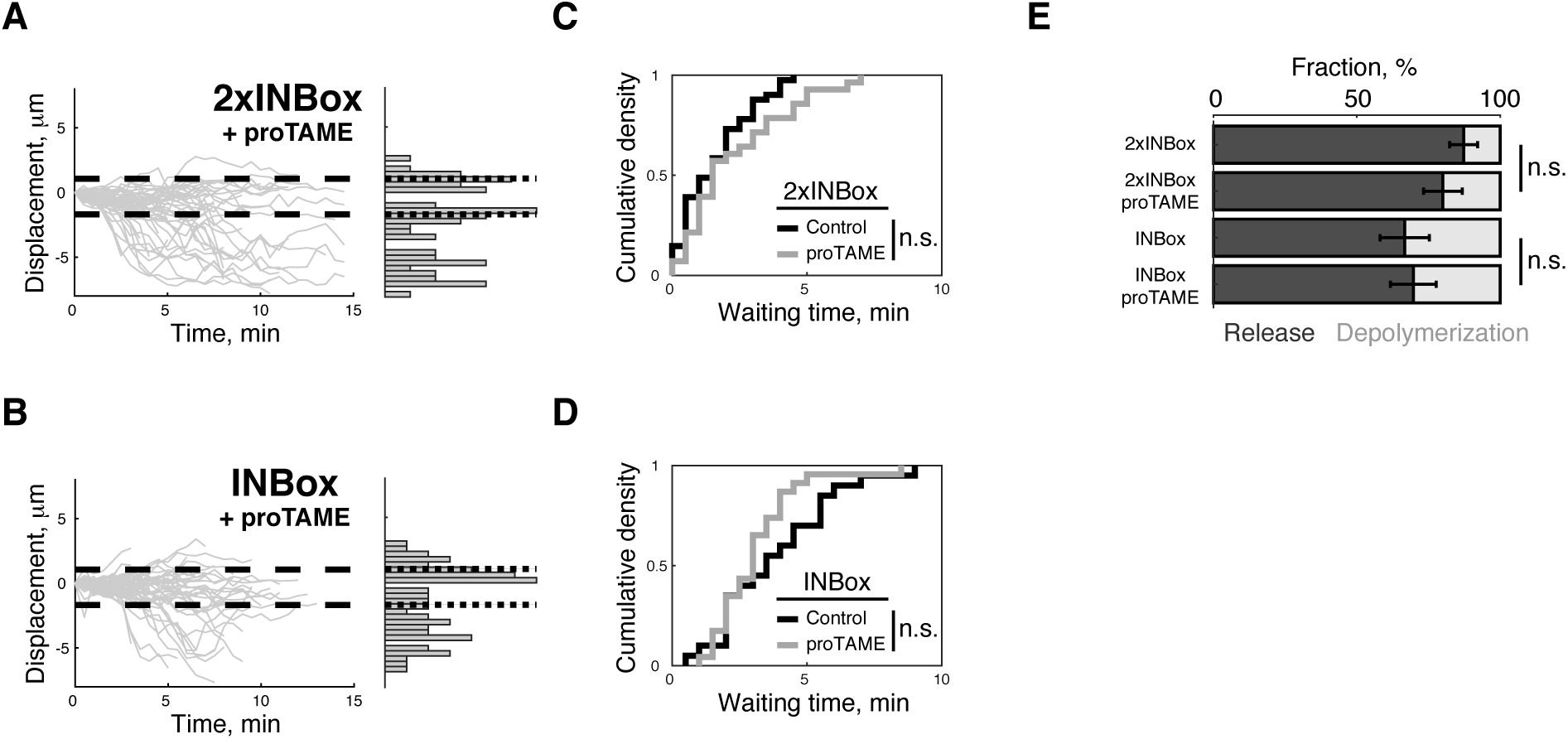
Microtubule release induced by Aurora B recruitment does not depend on time in mitosis. (**A-B**) Displacement of the activated kinetochore, plotted as in **Figure 2C**, with 20 μM proTAME to delay mitotic exit (2xINBox: n = 54; INBox: n = 56). (**C-D**) Waiting time of the released kinetochores after recruiting 2xINBox (control: median = 1.5 min, n = 41; proTAME: 1.5 min, n = 28) or INBox (control: 3.5 min, n = 20; proTAME: 3.0 min, n = 23). (**E**) The fraction of release vs. depolymerization events after recruiting 2xINBox or INBox. Differences between control and proTAME (C-E) are not statistically significant (P > 0.3).

**Figure S5.**
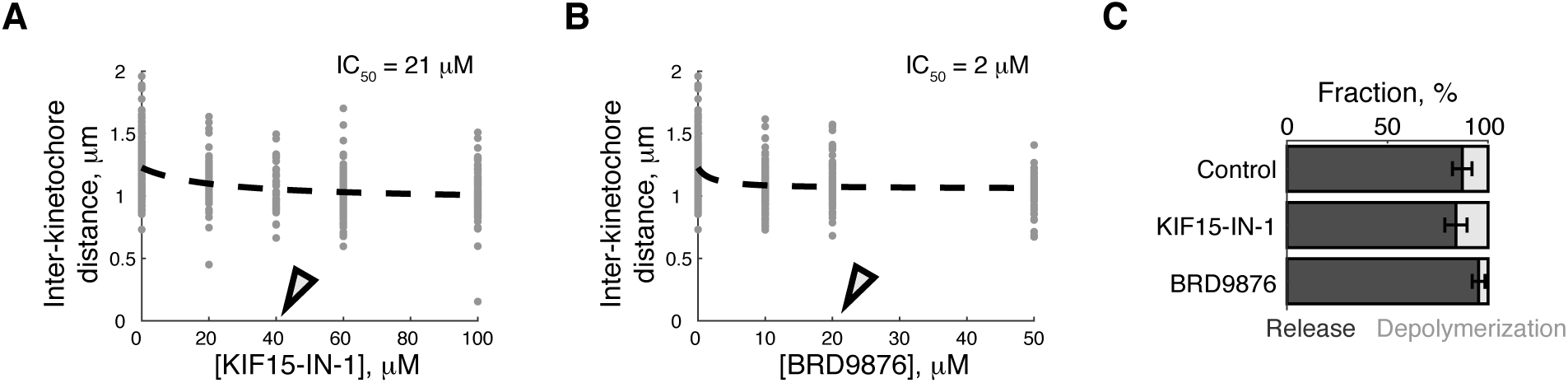
**Related to Figure 4**. (**A-B**) Inter-kinetochore distances at varying concentrations of rigor inhibitors for KIF15 (KIF15-IN-1: IC_50_ = 21 μM) or Eg5 (BRD9876: IC_50_ = 2 μM). Open triangles represent the concentrations used for 2xINBox recruitment assays in **Figure 4B-D** (KIF15-IN-1: 40 μM; KIF15-IN-1: 20 μM). (**C**) The fraction of release vs. depolymerization events from **Figure 4C-D** (Control: 87 ± 5% mean ± SEM, n = 47; KIF15-IN-1: 84 ± 6%, n = 44; BRD9876: 95 ± 3%, n = 43).

**Movie S1**.

**Related to Figure 1D**. INBox recruitment triggers poleward chromosome movement on a monopolar spindle. Left: 3xHalo-GFP-SPC25, right: mCherry-INBox-eDHFR.

**Movie S2**.

**Related to Figure 1E**. MCAK recruitment triggers poleward chromosome movement on a monopolar spindle. Left: 3xHalo-GFP-SPC25, right: MCAK-mCherry-eDHFR.

**Movie S3**.

**Related to Figure S2A**. Void-recruitment defines naturally-occurring chromosome oscillations on a bipolar spindle. Left: 3xHalo-GFP-SPC25, right: mCherry-eDHFR.

**Movie S4**.

**Related to Figure 2B**. INBox recruitment triggers microtubule release on a bipolar spindle. Left: 3xHalo-GFP-SPC25, right: mCherry-INBox-eDHFR.

**Movie S5**.

**Related to Figure S2C**. Example of INBox recruitment triggering microtubule depolymerization on a bipolar spindle. Left: 3xHalo-GFP-SPC25, right: mCherry-INBox-eDHFR.

**Movie S6**.

**Related to Figure 2C**. 2xINBox recruitment triggers microtubule release on a bipolar spindle. Left: 3xHalo-GFP-SPC25, right: INBox-mCherry-INBox-eDHFR.

## References and Notes

1. R. B. Nicklas, How cells get the right chromosomes. Science. 275, 632–7 (1997).

2. K. K. Sarangapani, C. L. Asbury, Catch and release: how do kinetochores hook the right microtubules during mitosis? Trends Genet. 30, 150–159 (2014).

3. M. A. Lampson, E. L. Grishchuk, Mechanisms to Avoid and Correct Erroneous Kinetochore-Microtubule Attachments. Biol. 6, 1 (2017).

4. S. Mukherjee et al., A Gradient in Metaphase Tension Leads to a Scaled Cellular Response in Mitosis. Dev. Cell. 49, 63-76.e10 (2019).

5. M. A. Lampson, I. M. Cheeseman, Sensing centromere tension: Aurora B and the regulation of kinetochore function. Trends Cell Biol. 21, 133–140 (2011).

6. M. Carmena, M. Wheelock, H. Funabiki, W. C. Earnshaw, The chromosomal passenger complex (CPC): From easy rider to the godfather of mitosis. Nat. Rev. Mol. Cell Biol. 13, 789–803 (2012).

7. V. Krenn, A. Musacchio, The Aurora B kinase in chromosome bi-orientation and spindle checkpoint signaling. Front. Oncol. 5, 225 (2015).

8. D. Liu, G. Vader, M. J. M. Vromans, M. A. Lampson, S. M. A. Lens, Sensing Chromosome Bi-Orientation Kinase from Kinetochore Substrates. Science. 323, 1350–1353 (2009).

9. T. Y. Yoo et al., Measuring NDC80 binding reveals the molecular basis of tension-dependent kinetochore-microtubule attachments. Elife. 7, 1–34 (2018).

10. L. J. García-Rodríguez, T. Kasciukovic, V. Denninger, T. U. Tanaka, Aurora B-INCENP Localization at Centromeres/Inner Kinetochores Is Required for Chromosome Bi-orientation in Budding Yeast. Curr. Biol. 29, 1536-1544.e4 (2019).

11. A. V. Zaytsev et al., Bistability of a coupled aurora B kinase-phosphatase system in cell division. Elife. 5, e10644 (2016).

12. Y. Asai et al., Aurora B kinase activity is regulated by SET/TAF1 on Sgo2 at the inner centromere. J. Cell Biol. 218, 3223–3236 (2019).

13. C. S. Campbell, A. Desai, Tension sensing by Aurora B kinase is independent of survivin-based centromere localization. Nature. 497, 118–121 (2013).

14. A. J. Broad, K. F. DeLuca, J. G. DeLuca, Aurora B kinase is recruited to multiple discrete kinetochore and centromere regions in human cells. J. Cell Biol. 219, e201905144 (2020).

15. A. J. Broad, J. G. Deluca, The right place at the right time: Aurora B kinase localization to centromeres and kinetochores. Essays Biochem., EBC20190081 (2020).

16. M. A. Lampson, K. Renduchitala, A. Khodjakov, T. M. Kapoor, Correcting improper chromosome –spindle attachments during cell division. Nat. Cell Biol. 6, 232–237 (2004).

17. A. A. Ye et al., Aurora A Kinase Contributes to a Pole-Based Error Correction Pathway. Curr. Biol. 25, 1842–1851 (2015).

18. L. Chmátal, K. Yang, R. M. Schultz, M. A. Lampson, Spatial Regulation of Kinetochore Microtubule Attachments by Destabilization at Spindle Poles in Meiosis i. Curr. Biol. 25, 1835–1841 (2015).

19. E. R. Ballister, C. Aonbangkhen, A. M. Mayo, M. A. Lampson, D. M. Chenoweth, Localized light-induced protein dimerization in living cells using a photocaged dimerizer. Nat. Commun. 5, 5475 (2014).

20. H. Zhang et al., Optogenetic control of kinetochore function. Nat. Chem. Biol. 13, 1096–1101 (2017).

21. E. J. Banigan et al., Minimal model for collective kinetochore-microtubule dynamics. Proc. Natl. Acad. Sci. 112, 12699–12704 (2015).

22. T. M. Kapoor, T. U. Mayer, M. L. Coughlin, T. J. Mitchison, Probing spindle assembly mechanisms with monastrol, a small molecule inhibitor of the mitotic kinesin, Eg5. J. Cell Biol. 150, 975–988 (2000).

23. C. E. Walczak, S. Gayek, R. Ohi, Microtubule-Depolymerizing Kinesins. Annu. Rev. Cell Dev. Biol. 29, 417–441 (2013).

24. V. Sikirzhytski et al., Direct kinetochore-spindle pole connections are not required for chromosome segregation. J. Cell Biol. 206, 231–243 (2014).

25. M. W. Elting, M. Prakash, D. B. Udy, S. Dumont, Mapping Load-Bearing in the Mammalian Spindle Reveals Local Kinetochore Fiber Anchorage that Provides Mechanical Isolation and Redundancy. Curr. Biol. 27, 2112-2122.e5 (2017).

26. B. Polak, P. Risteski, S. Lesjak, I. M. Tolic, PRC 1-labeled microtubule bundles and kinetochore pairs show one-to-one association in metaphase. EMBO Rep. 18, 217–230 (2017).

27. L. C. Kapitein et al., The bipolar mitotic kinesin Eg5 moves on both microtubules that it crosslinks. Nature. 435, 114–8 (2005).

28. Y. Shimamoto, S. Forth, T. M. Kapoor, Measuring Pushing and Braking Forces Generated by Ensembles of Kinesin-5 Crosslinking Two Microtubules. Dev. Cell. 34, 669–681 (2015).

29. M. Uteng, C. Hentrich, K. Miura, P. Bieling, T. Surrey, Poleward transport of Eg5 by dynein-dynactin in Xenopus laevis egg extract spindles. J. Cell Biol. 182, 715–726 (2008).

30. E. G. Sturgill, R. Ohi, Kinesin-12 differentially affects spindle assembly depending on its microtubule substrate. Curr. Biol. 23, 1280–1290 (2013).

31. H. Drechsler, T. McHugh, M. R. Singleton, N. J. Carter, A. D. McAinsh, The Kinesin-12 Kif15 is a processive track-switching tetramer. Elife. 3, e01724 (2014).

32. D. J. Needleman et al., Fast Microtubule Dynamics in Meiotic Spindles Measured by Single Molecule Imaging: Evidence That the Spindle Environment Does Not Stabilize Microtubules. Mol. Biol. Cell. 21, 323–333 (2010).

33. M. E. Dumas et al., Dual inhibition of Kif15 by oxindole and quinazolinedione chemical probes. Bioorganic Med. Chem. Lett. 29, 148–154 (2019).

34. G.-Y. Chen et al., Eg5 Inhibitors Have Contrasting Effects on Microtubule Stability and Metaphase Spindle Integrity. ACS Chem. Biol. 12, 1038–1046 (2017).

35. A. Peña, A. Sweeney, A. D. Cook, M. Topf, C. A. Moores, Structure of Microtubule-Trapped Human Kinesin-5 and Its Mechanism of Inhibition Revealed Using Cryoelectron Microscopy. Structure. 28, 450–457 (2020).

36. B. Akiyoshi et al., Tension directly stabilizes reconstituted kinetochore-microtubule attachments. Nature. 468, 576–579 (2010).

37. N. T. Umbreit et al., The Ndc80 kinetochore complex directly modulates microtubule dynamics. Proc. Natl. Acad. Sci. 109, 16113–16118 (2012).

38. K. K. Sarangapani, B. Akiyoshi, N. M. Duggan, S. Biggins, C. L. Asbury, Phosphoregulation promotes release of kinetochores from dynamic microtubules via multiple mechanisms. Proc. Natl. Acad. Sci. 110, 7282–7287 (2013).

39. D. Cimini, X. Wan, C. B. Hirel, E. D. Salmon, Aurora Kinase Promotes Turnover of Kinetochore Microtubules to Reduce Chromosome Segregation Errors. Curr. Biol. 16, 1711–1718 (2006).

40. S. F. Bakhoum, S. L. Thompson, A. L. Manning, D. A. Compton, Genome stability is ensured by temporal control of kinetochore-microtubule dynamics. Nat. Cell Biol. 11, 27–35 (2009).

41. J. P. I. Welburn et al., Aurora B Phosphorylates Spatially Distinct Targets to Differentially Regulate the Kinetochore-Microtubule Interface. Mol. Cell. 38, 383–392 (2010).

42. T. M. Kapoor et al., Chromosomes can congress to the metaphase plate before biorientation. Science. 311, 388–91 (2006).

43. A. L. Knowlton, W. Lan, P. T. Stukenberg, Aurora B Is Enriched at Merotelic Attachment Sites, Where It Regulates MCAK. Curr. Biol. 16, 1705–1710 (2006).

44. K. J. Salimian et al., Feedback control in sensing chromosome biorientation by the aurora B kinase. Curr. Biol. 21, 1158–1165 (2011).

45. P. Trivedi et al., The binding of Borealin to microtubules underlies a tension independent kinetochore-microtubule error correction pathway. Nat. Commun. 10, 682 (2019).

46. A. V. Zaytsev, E. L. Grishchuk, Basic mechanism for biorientation of mitotic chromosomes is provided by the kinetochore geometry and indiscriminate turnover of kinetochore microtubules. Mol. Biol. Cell. 26, 3985–3998 (2015).

47. N. J. Ganem, D. A. Compton, Functional Roles of Poleward Microtubule Flux During Mitosis. Cell Cycle. 5, 481–485 (2006).

48. K. A. Knouse, K. E. Lopez, M. Bachofner, A. Amon, Chromosome Segregation Fidelity in Epithelia Requires Tissue Architecture. Cell. 175, 200–11 (2018).

49. Z. Lin et al., TTC5 mediates autoregulation of tubulin via mRNA degradation. Science. 367, 100–104 (2020).

50. A. K. Gillingham, S. Munro, The PACT domain, a conserved centrosomal targeting motif in the coiled-coil proteins AKAP450 and pericentrin. EMBO Rep. 1, 524–529 (2000).

51. H. Zhang, D. M. Chenoweth, M. A. Lampson, Optogenetic control of mitosis with photocaged chemical dimerizers (Elsevier Inc., ed. 1, 2018), vol. 144.

52. P. Khandelia, K. Yap, E. V. Makeyev, Streamlined platform for short hairpin RNA interference and transgenesis in cultured mammalian cells. Proc. Natl. Acad. Sci. 108, 12799–12804 (2011).

53. T. Maney, A. W. Hunter, M. Wagenbach, L. Wordeman, Mitotic centromere-associated kinesin is important for anaphase chromosome segregation. J. Cell Biol. 142, 787–801 (1998).

54. S. K. Talapatra, B. Harker, J. P. I. Welburn, The C-terminal region of the motor protein MCAK controls its structure and activity through a conformational switch. Elife. 2015, 1–55 (2015).

55. D. A. Skoufias et al., S-trityl-L-cysteine is a reversible, tight binding inhibitor of the human kinesin Eg5 that specifically blocks mitotic progression. J. Biol. Chem. 281, 17559–17569 (2006).

56. X. Zeng et al., Pharmacologic inhibition of the anaphase-promoting complex induces a spindle checkpoint-dependent mitotic arrest in the absence of spindle damage. Cancer Cell. 18, 382–395 (2010).

57. Rieder, Conly L., G. Cassels, Correlative Light and Electron Microscopy of Mitotic Cells in Monolayer Cultures. Methods Cell Biol. 61, 297–315 (1999).

